# Repetition Dissociates Pointer and Content-based Representations in Visual Working Memory: Contrasting the CDA with Multivariate Shape Decoding

**DOI:** 10.64898/2026.06.28.735064

**Authors:** Dock H. Duncan, Güven Kandemir, Christian N. L. Olivers

## Abstract

Memorizing a new phone number or address is hard at first, but becomes easier with repetition, as information shifts from working memory to long-term memory. Here we investigated how repetition affects the storage and transition of different aspects of mnemonic information by comparing univariate neural markers of active object storage with multivariate decoding of memory content. Thirty participants encoded lateralized stimuli from a continuous shape space into memory. Memory items were repeated six times in a row to induce learning. In line with earlier work, EEG recordings revealed that repetition led to a reduction in contralateral delay activity (CDA), a measure of active storage that has been taken to reflect a pointer-like representation of the individual object or its original source. In contrast, shape decoding during the retention and also after an impulse perturbation remained constant across repetitions. These results suggest that learning over repetitions reflects the abolishment of active and individuated object memory representations while passive, source-independent memory representations are retained.

**Highlights:**

- We recorded EEG as participants maintained a shape item, drawn form a continuous shape space, in working memory. These memory items repeated six times in a row before they changed to a new item.
- We replicated the well-known CDA decrement effect from repetition, a process thought to index participants’ shift from relying on working memory to long term memory.
- Multivariate decoding of shape identity was stable across repetitions, neither decreasing nor reliably increasing with repetition. This was true both after encoding and also following a task irrelevant ‘ping’.
- These results dissociate the popular CDA ERP, a measure of memory load, from actual working memory content decodable via multivariate approaches.

## 1. Intro

Working memory is a fundamental cognitive ability allowing for the flexible retention and manipulation of information, thus serving adaptive behavior (Baddeley, 2020; Cowan et al., 2005; Engle et al., 1999; Unsworth et al., 2014). Moreover, working memory serves as an important gateway for long-term memory, as in particular the repetition of information in working memory enhances learning, thus promoting more accurate and durable memory performance.

Within the visual domain, the repetition-induced transition from visual working memory (VWM) to long term memory has found empirical support in changes in the contralateral delay activity (CDA), an event-related component in the electroencephalogram (EEG). The CDA is a sustained negative voltage signal emerging contralaterally during the delay period of a VWM task, and is known to be sensitive to memory load (Vogel & Machizawa, 2004; see Balaban & Luria, 2025 for a review). Importantly, Carlisle et al. (2011) observed that the CDA decreases with repeated exposure to the same memorandum across seven trials in a row. Carlisle and colleagues proposed that this CDA decrease marks a shift in mnemonic representations from primarily relying on working memory to relying on long term memory. With learning, the object no longer needs to be actively represented in VWM, and thus any load-related index should diminish. Follow-up work has replicated and expanded the observation of a CDA decrease with repetition (Drew et al., 2018; Gunseli, Olivers, et al., 2014; Heinen et al., 2022; Özdemir et al., 2024; Woodman et al., 2013).

An open question is: what changes in VWM across repetitions, and what the CDA then ultimately reflects in this process. We focus on two options here, the first of which is that the CDA reflects the activity of the memory content itself. While the CDA itself is a coarse univariate measure that is not known to contain content information as such, it may nevertheless reflect the aggregated strength of stimulus-specific activity of feature-sensitive neurons. With learning, this activity-based representation in VWM then transitions to a passive, connectivity-based long-term representation, resulting in a reduced CDA. The second possibility is that instead of representing aggregated content activity, the CDA provides important episodic, or contextual, binding of the memory content. Specifically, the CDA has been argued to reflect a “pointer”, “object file”, or “tokenization” process, in which the spatiotemporal context of the memorized object is being maintained in order to enable the individuation of separate instances of identical objects, as well as provide useful retrieval cues (Awh & Vogel, 2025; Balaban & Luria, 2025). In line with such a content-independent pointer system, the CDA amplitude appears to be tied to the number of objects in memory and not the number of features (Balaban et al., 2018, 2019; Balaban & Luria, 2015, 2016b, 2016a; Diaz et al., 2021; Luria & Vogel, 2014; Woodman & Vogel, 2008). Under this perspective, a repetition-induced decline in CDA would reflect a decrease in pointer activity, rather than content. In fact, there is good reason to assume that the active maintenance of the contextual information diminishes. First and foremost, the repetition-induced decline of the CDA has been studied in tasks in which the spatial (and temporal) source of the memory object is irrelevant to the memory test at the end of each trial, which is always about memory content with no relation to encoding location. Hence, while spatial location might initially provide a useful retrieval cue towards the content, with learning, the added value of also remembering the original location diminishes. This may be further aggravated by the fact that this location changes randomly between trials, thus making location an increasingly less specific retrieval cue. In sum, the spatial origin of the memory simply becomes less useful with repetition, and if the CDA indeed reflects that spatial origin, it should diminish with time.

If the repetition-induced CDA reduction indeed merely reflects a reduction of pointer information in VWM, the question arises what repetition does to the feature content. Will the stimulus-specific information also diminish with learning, or will it behave differently? If the latter is true, then repetition-based learning provides another handle on dissociating content-specific and content-general processes in VWM. To this end we adopted a multivariate decoding approach which also relies on neuronal activity, and has been demonstrated to be sensitive to stimulus-specific memory content such as orientation and color (in fMRI: Brouwer & Heeger, 2009; Harrison & Tong, 2009; Kamitani & Tong, 2005; as well as in EEG: Hajonides et al., 2021; Wolff et al., 2015). Here we chose to draw stimuli from a recently developed continuous shape space (Li et al., 2020), allowing us to observe whether successful decoding of stimulus-specific content from the EEG can be extended to shape space (see Figure 1A for an illustration of this space). Participants were presented with two lateralized shape stimuli at the beginning of each trial and asked to encode one into their memory (target indicated via color; Figure 1B). Following a 1,000 ms retention interval, a salient task-irrelevant impulse was presented for 100 ms followed by a further retention interval of 500 ms. Participants were then presented with a probed at fixation that either matched their encoded memory item, or was a foil drawn from nearby in the stimulus space. Crucially, the same memory target was repeated for six trials in a row (following Carlisle et al., 2011; Woodman et al., 2013). To investigate the consequences of repetition on VWM, in addition to measuring any changes in CDA, we sought to assess how repetition affects multivariate content-based signals. We anticipated three potential outcomes (all preregistered): Firstly, shape decoding during the delay could decrease with repetition, thus matching our expected CDA decrease. This would imply that the stimulus-specific signals themselves are no longer represented in neural activity, consistent with a shift towards more passive and potentially more durable connectivity-based representations. If so, this outcome would provide some, though inconclusive, evidence that the CDA and content-based signals may in the end reflect the same activity. Secondly, while the CDA decreases, decoding performance may remain constant with repetition, indicating little change in content-related activity across repetitions. If so then this would indicate a dissociation between the CDA and content-related activity across repetitions, reflecting a decline in pointer information while content is retained. Thirdly, shape decoding during the delay may increase with repetition. This too would provide clear evidence for a dissociation between the CDA and content-based signals. Moreover, it would suggest that repetition actually strengthens VWM representations rather than offloading them to LTM – at least within the range of repetitions typical for this paradigm.

**Figure 1.**
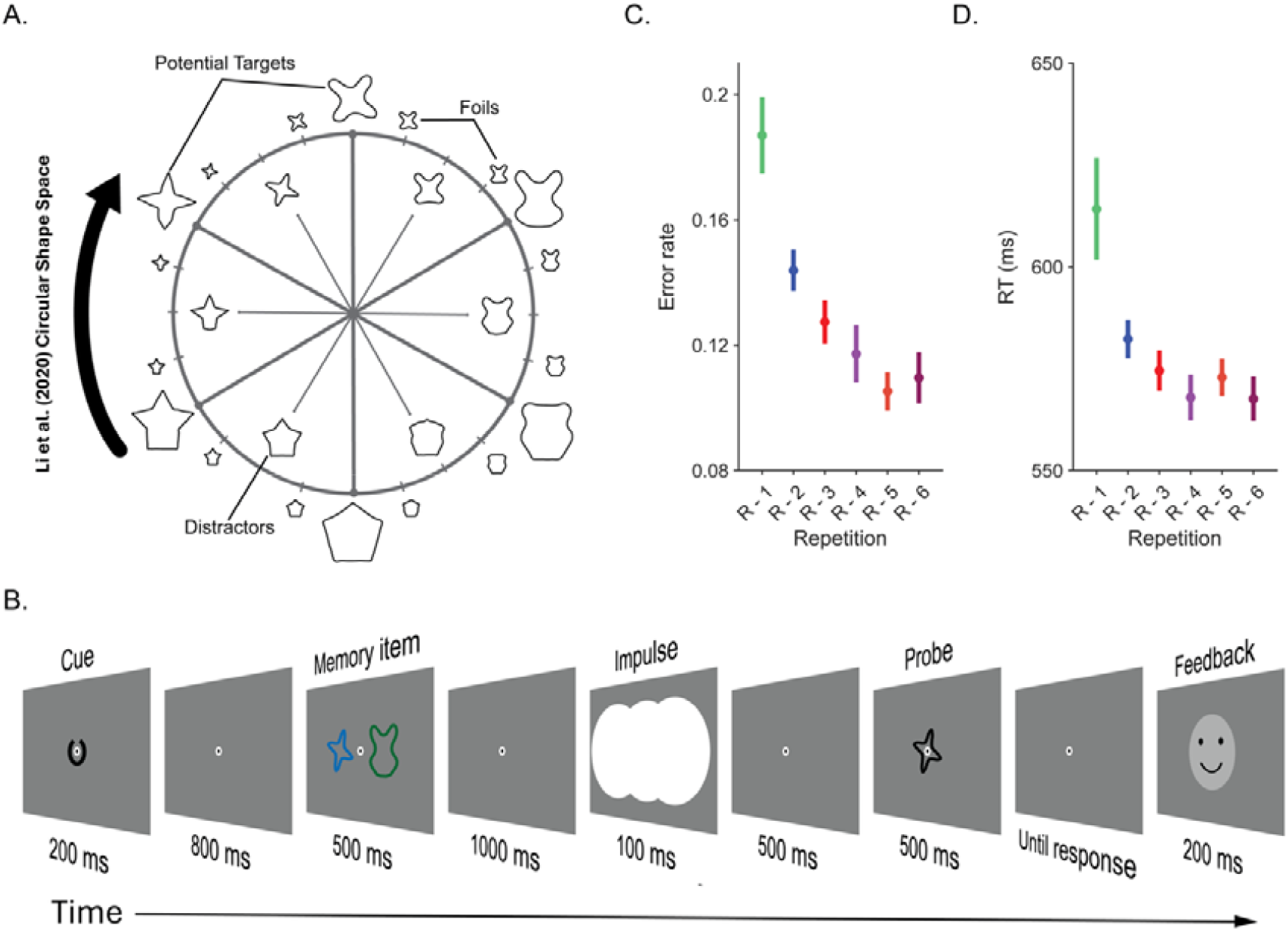
Stimuli and trial flow plus behavioral data. **Note**: **(A)** visualization of the validated circular shape space from Li et al. (2020). Highlighted are the six memory items used throughout the experiment (large and present on the outside of the circle). Beside these potential target shapes, 15° to their right and left in the visualization were the foils which could be presented during test phase. Note that foils were never another potential memory item. Inside the circle are presented the six nontarget distractors that would also be presented during the encoding window. These nontarget shapes were also never other potential target shapes. Note that all stimuli are presented at their approximate position within the 360° stimulus space. **(B)** Demonstration of a single trial. In each trial, the cue is rotated by 60° in clockwise direction to indicate the repetition count (in this case, shows first presentation or Repetition 1. A memory item (in this case the shape in blue), presented alongside a distractor (in this case, shape in green), is remembered throughout a delay period to make a same/different judgement on a probe (a shape presented centrally in black). Impulse stimuli (the task irrelevant white disks) were presented during delay to improve EEG decoding and subjects were required to press ‘C’ or ‘M’ to respond based on subject wise response-mapping. **(C)** Changes in response time across repetition conditions. **(D)** Changes in accuracy across repetition conditions.

VWM activity may not only wane across trials, but also within trials, as the CDA but also decoding-based signals tend to weaken with increasing delay. This, in and of itself, may reflect an offloading process where a more passive storage system is recruited (Mongillo et al., 2008; Polyn & Woodman, 2025; Stokes, 2015). In order to study the effect of repetition later in the delay intervals we adopted the impulse perturbation technique (often referred to as ‘pinging’; Stokes et al., 2013; Wolff et al., 2015, 2017) wherein a salient, task irrelevant visual stimulus is presented during the retention interval during a period when multivariate decoding of memory items often drops to baseline. The idea is that by then the memory has been stored in connectivity rather than activity, and the stimulus-induced perturbation then evokes broadband neural activity. which is sent through the memory network, allowing for the memorized item to be decoded. Here we assessed if repetition resulted in a stronger reliance on such activity-silent representations. If so, this would suggest that these activity-silent representations as have been observed during working memory intervals within a trial, also underlie longer-term learning.

## 1. Methods

The study was preregistered at https://osf.io/hxdfc. The data, analysis scripts and analysis outputs are publicly available here: https://osf.io/gupzq

### 2.1 Participants

Thirty young adults volunteered to participate (24 female, M_Age_ = 23.7, s.d._Age_ = 4.30) in return for monetary reward (50 €). To acquire adequate power in impulse driven decoding, the sample size was the same as earlier studies (Kandemir, Wilhelm, et al., 2024; Kandemir & Akyürek, 2023; Kandemir & Olivers, 2024). A verbal and a written explanation of the experimental procedure, as well as the aim of the project was provided, and participants signed a written consent form declaring their compliance with data storage practices and agreement to participation. The experimental protocol was approved by the Scientific and Ethics Review Board of the Faculty of Behavioral and Movement Sciences at VU University Amsterdam and adhered to the declaration of Helsinki. The rationale and the methodology of the study were preregistered (https://osf.io/hxdfc).

### 2.2 Stimuli and Apparatus

The display background in the experiment was gray (RGB = 128, 128, 128). A fixation dot formed by a white dot (0.25 degrees of visual angle in diameter [d.v.a]) surrounded by a black circle (0.3 d.v.a. diameter) was displayed at the center of the screen throughout a trial.

The sample of memory items, distractors and probes were non-filled 2-dimensional shape figures sampled from a continuous circular shape space (Li et al., 2020). For memory items, 6 shapes with 60 degrees angular distance were selected (40°, 100°, 160°, 220°, 280°, 340°; Figure 1A), whereas distractors were 30 degrees apart, also with 60 degrees angular distance (10°, 70°, 130°, 190°, 250°, 310°). The twelve false probes were shapes +/-15° away from each memory item. The memory items, distractors and the probe were 6.5 degrees visual angle (d.v.a.) in width and height. The target and the distractor were presented in blue (RGB= 0, 106, 176) and aqua (RGB= 0, 102, 51) and in every 432 trials the colors were swapped, whereas the probe was always in black (RGB = 0, 0, 0).

In each trial, memory items were preceded by a cue, indicating the repetition count for the target item. The cue was a black (RGB = 0, 0, 0), non-closed circle (Landolt C, 1.625 d.v.a. diameter) with a 10 ° arc-opening displayed at the center of the screen, surrounding the fixation dot. The opening was directed to 12 o’clock to indicate the first item in sequence and it was rotated in clockwise direction by 60 ° in each subsequent presentation. The impulse stimulus consisting of 3 overlapping white disc (RGB= 255, 255, 255) with a diameter of 13 d.v.a., centered at the middle of the screen as well as centered at each item location at 4 d.v.a. away from the origin of the screen.

The stimuli were created and the experimental flow was controlled by Psychtoolbox extension for MATLAB (Brainard & Vision, 1997; Kleiner et al., 2007). The output was a 23.8 inch (60,452 cm on diagonal) ASUS ROG Strix XG248Q monitor with a resolution 1920 by 1080 pixels and a refresh rate of 240 Hz. A regular keyboard was used to collect the responses where buttons ‘C’ and ‘M’ were pressed to indicate whether the probe was ‘the same’ or ‘different’ relative to memory item in each trial. The response mappings alternated across each experimental session, counterbalanced across participants. A desktop-mounted Eyelink 1000 Plus (SR-Research) monitored and recorded gaze position and other eye data at 1000 Hz sampling rate.

### 2.3 Procedure

Each participant first received information about the experimental procedure and signed the consent form. Next, sitting in a dimly lit room, a practice block with 48 trials identical to experimental trials were completed to warm up for the experiment.

Experiment blocks were self-initiated by the participant and began with the message ‘Get Ready’ presented for 400 ms. This was followed by a blank screen lasting 600 ms and then the cue was presented for 200 ms at the center of the screen. After another delay for 800 ms, a memory item and a distractor were presented simultaneously for 500 ms, each centered at 4 d.v.a. lateral to the left and right of the center of the screen. After another delay with 1,000 ms duration, the impulse stimulus was presented for 100 ms and the final delay for 500 ms. Finally the probe that was either the same shape as the memory item or a shape -/+ 15° to the relevant memory item was presented for 500 ms. The trial stopped when participants indicated whether the probe was ‘the same’ or ‘different’ by pressing ‘C’ or ‘M’ on the keyboard. The pairing of response with buttons switched between experimental sessions, with its order counterbalanced across participants. 100 ms after response, a feedback stimulus consisting of a happy or sad smiley indicating the accuracy of the response was presented for 200 ms, After a delay with random jitter ranging from 200 – 500 ms, the next trial began automatically. For a visualization of the complete trial flow, see Figure 1B The study contained four sessions each with 432 trials per session. Each session was further made up of 18 blocks with 24 trials per block. Within a block each target shape could only be target once, meaning that repetition count did not exceed 6 within a block.

Participants could take self-paced breaks between blocks as well as between sessions to combat fatigue. In these breaks participants were informed of their accuracy and reaction times across all blocks in that session so far. In total, each participant completed 1728 trials distributed over 4 consecutive sessions that lasted ∼45 minutes each. All practice and test trials were completed on the same day.

### 2.4 EEG Acquisition and Preprocessing

The EEG was recorded at 1024 Hz from 64 channels arranged according to international 10-20 system via BioSemi Active 2 system. The reference was the average of two electrodes placed on mastoids. Bipolar EOG was recorded from electrodes placed below and above the right eye (Vertical) and lateral to the external canthi (Horizontal).

Offline, the data was re-referenced and downsampled to 500 Hz. Next, it was bandpass filtered (0.1 Hz highpass – 40 Hz lowpass). The Matlab toolboxes Fieldtrip (Oostenveld et al., 2011) and EEGLab (Delorme & Makeig, 2004) were used for the preprocessing, filtering and the visualization of the EEG data.

For ICA, a copy of data was separately downsampled to 1-40 Hz and ICA was performed by the *runica* function from EEGLab. The main data for each subject was corrected for blink induced artefacts individually by removing ICA components selected via visual inspection after the visualization of ICA maps. Post ICA, data was visually inspected for artefacts and on average, 0.73 noisy channels were replaced by spherical spline interpolation. The data was separately epoched relative to cue and impulse stimulus onsets (- 200– 1000 ms relative to cue and -200 – 500 ms relative to impulse). All epochs were inspected for artefacts visually and effected trials were marked for exclusion from analyses.

### 2.5 EEG Analyses

#### 2.5.1 Contralateral delay activity

For the Contralateral delay activity (CDA) measure, we used pre-stimulus baseline (-200 to 0 ms relative to memory item onset), at which mean activity in each electrode was calculated and subtracted from the remainder of the electrode measure, just as previous studies (Carlisle et al., 2011; Özdemir et al., 2024; Vogel & Machizawa, 2004). Next, the data were first grouped according to each repetition condition. In a second analysis, repetitions 1 & 2, 3 & 4, and 5 & 6 were grouped together to match the trial pairing in our multivariate analysis (see below). During this grouping, we ensured all shapes and locations were equally represented within the new repetition condition by randomly sub sampling trials. In order to avoid selection effects, this procedure was repeated 100 times using random subsampling. The CDA was measured as the difference of signal in the contralateral electrode from ipsilateral electrode based on the target item location using electrodes PO7 and PO8. The difference wave was smoothened using Gaussian filter (kernel = 2 time points, 16 ms). To contrast across conditions, we calculated mean difference wave once relative to memory item offset from 500 to 1500 ms.

### 2.5.2 Decoding Analysis

Consistent with our preregistration, decoding analysis was first conducted on all trials in order to test discrimination of shape stimuli, then the same analysis was conducted on a subset of trials grouped into distinct bins reflecting different repetition conditions. The aim was to increase the power as the successful decoding of VWM feature content from EEG tends to require a high density of trials (Kandemir, Wilhelm, et al., 2024; Kandemir & Olivers, 2026). As such, trials in repetitions 1 and 2, 3 and 4, and 5 and 6 were grouped together while trial count, memory item type and presentation location was equalized across bins via random sampling.

All multivariate analyses were completed using EEG data from 17 posterior electrodes (PO7, PO3, POz, Pz, P1, P3, P5, P7, Oz, P2, P4, P6, P8, O1, PO8, PO4, O2), based on earlier studies using impulse-driven decoding (Wolff et al., 2017; Wolff, Jochim, et al., 2020). Decoding was completed using an 8-fold cross-validation technique: trials were first distributed into 8 groups with stratified sampling, so that each condition would be equivalently represented in each group. Next, 7 of these folds were used as training set, as these trials were distributed to six bins, each for the respective target shape in that trial. The number of trials in each bin was equalized by randomly sampling as many trials as the bin with least trial count. The EEG data in each bin was averaged to form a shape-specific ERP pattern. A custom shrinkage estimator (Ledoit & Wolf, 2003) was used to compute the covariance matrix by using all trials in the training set. Each trial in the remaining test set was compared to these patterns and dissimilarity was measured in Mahalanobis distance (De Maesschalck et al., 2000). For each time-point in each trial, six distance measures were calculated. Distance measures were ordered by centering the target, mean-centered and the sign was flipped to reflect similarity across conditions. Mean of cosine convoluted, sign-reversed Mahalanobis distances yielded decoding accuracy at each time point. To reduce variation in decoding accuracy due to differences in training set, each decoding analyses was repeated 100 times and the product was the average across repetitions and trials, which was smoothened using a Gaussian filter over the time dimension (kernel = 2 time points, 16 ms). Data from all participants was then tested for statistical significance with permutation tests. In order to assess the impact of repetition on decoding, we conducted decoding analyses separately for each repetition condition and the trial-count was equalized across all shapes and repetition conditions. Due to the presence of a CDA in the window immediately before impulse presentation, and because this CDA scaled precisely with repetition (which was our main factor of interest) we were unable to use a conventional baseline in the window immediately before impulse perturbation. We therefore we applied a moving baseline and focused on the residual dynamic signal (Kandemir, Wolff, et al., 2024; Wolff et al., 2019). For this, data on each channel within a 100 ms wide window was first subtracted and the residual activity was down then sampled to 100 Hz, yielding 10 time-points in each window. This data was concatenated across all channels, providing 170 data features at each time-point which were then used in our impulse driven multivariate analyses. This procedure was repeated for each time point by sliding the window to the following 100 ms window, thereby isolating evoked changes in the EEG signal from the impulse without baselining over the fading CDA window (note that we did not specify this baselining procedure in our preregistration as we had not considered then that the CDA would interfere with a standard baselining approach).

We also controlled our analyses for the contribution of eye movements on the shape decoding (i.e. Mostert et al., 2018). For this, we first baselined (x,y) coordinates extracted from eye-tracker by subtracting -200 – 0 ms window prior to lateralized memory item presentation. Then we repeated the exact same decoding approach as above. The analysis was conducted separately for early CDA epoch and impulse epoch, though the same baseline was used for both.

### 2.5.3 Statistical assessment

The statistical significance of decoding was tested with non-parametric sign permutation test with cluster correction (Spaak, 2015). For this, the sign of each observation was flipped randomly with 0.5 probability and the statistic across observations was recomputed to generate a null distribution across 100,000 permutations. Timepoints in observed and permuted data exceeding a threshold (*p* < 0.05, two-tailed) were grouped into clusters. Observed cluster statistics were then compared to maximum cluster statistics from the null distribution and the probability of observing clusters of that magnitude is reported as the *p* value.

The impact of repetition on CDA was tested on the average across time window of interest. These values were first tested with an omnibus repeated measures analysis of variance (RM-ANOVA) and then individual conditions were contrasted with Bonferroni corrected t-tests.

## 2. Results

### 2.1 Behavior

Figures 1B and 1C show the average error and RT as a function of repetition. Performance was worst at the beginning of a sequence (Repetition 1) and gradually improved as the memory template was repeated. This was confirmed by a one-way repeated measure analysis of variance (ANOVA) taking error rate as the dependent variable and repetition number (1-6) as factors (*F*(2.90, 84.22)= 54.25, *p* < 0.001, η^2^ = 0.65; Figure 1C). Pairwise comparisons showed that this was mostly driven by the reliably higher error rates during the initial presentation (all p’s < 0.001, with Bonferroni correction) as well as Repetition 2 (all p’s *p* < 0.05, with Bonferroni correction). A similar picture emerged when analyzing reaction time, (*F*(1.77, 51.44) = 27.30, *p* < 0.001, η^2^ = 0.49; Figure 1D), with slowest responses observed during the first trials of the repetition sequence relative to all other repetitions (all p’s < 0.001, with Bonferroni correction).

### 2.2 CDA

Figure 2 shows the CDA as a function of repetition. We calculated the CDA as the difference in signal between electrodes contralateral and ipsilateral to the target (PO7/PO8; Figure 2A, see methods), which we then averaged across the 500 – 1500 ms time window (Figure 2B). A repeated-measures ANOVA on the CDA amplitude revealed a trend for the main effect of repetition, which was not significant after Greenhouse-Geisser correction (*F*(3.27, 94.81) = 2.33, *p* = 0.074). Visual inspection of Figure 2B indicates that the first presentation resulted in the CDA with the largest amplitude, which then declined for all subsequent repetitions.

**Figure 2.**
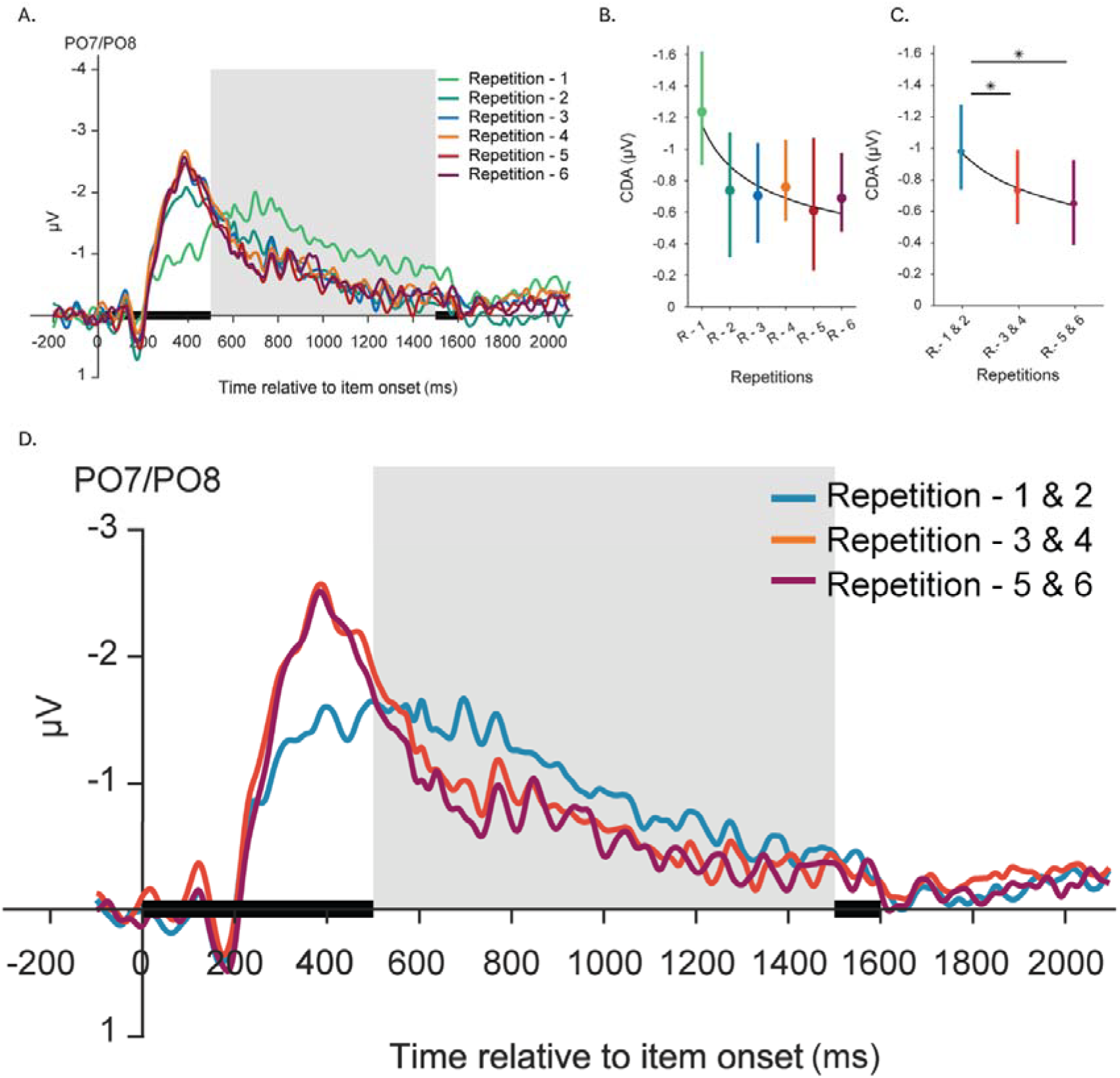
CDA Results. **Note**: **(A)** Contralateral delay activity per repetition as a function of time and **(B)** averaged over 500-1500 ms window over all trials with a fitted power function overlayed on the data. **(C)** Temporally averaged CDA within 500 – 1500 ms period when Repetition 1&2, 3&4 and 5&6 are pooled together and trial counts are equalized via subsampling. **(D)** Time course CDA over pooled repetitions. A & D: Solid colored lines show mean PO7/PO8 difference. Gray region marks 500-1500 ms window of interest for temporal averaging. B & C: Average CDA within window of interest where dot marks mean and bars indicate 95 % CI for each repetition condition. Asterisks indicates statistically significant difference between repetitions (*, p < 0.05; **, p < 0.01)

Consistent with our preregistration, we next pooled the CDA for presentations 1&2, 3&4 and 5&6 together while matching test side and shape type trial counts across bins so that our CDA condition matched the trial grouping that we used in our decoding analysis (see methods). As can be seen from Figures 2C and D, we now observed a reliable decline in CDA across repetitions (*F*(2. 58)= 5.10, *p*= 0.009, η^2^ = 0.15; Figures 2C). Pairwise comparisons revealed that the CDA for Repetition 1&2 was significantly larger than Repetition 3&4 (*t*(29) = 2.593, *p* = 0.044, η^2^ = 0.154) as well as Repetition 5&6 (*t*(29) = 2.88, *p* = 0.022, η^2^ = 0.171) whereas Repetition 3&4 did not differ from Repetition 5&6 (*t*(29) = 0. 30, *p* _*uncorrected*_ = 0.53). Finally, no CDA could be reliably observed after impulse perturbation. This stands in contrast to some work that has shown the CDA is resistant to interruption (Hakim et al., 2020) but aligns with other work suggesting task irrelevant perturbations can influence working memory behavior (Duncan & Van der Stigchel, 2020; Yang et al., 2025).

### 2.3 Decoding

Figure 3 shows shape decoding performance after stimulus presentation and after the impulse. Trial-specific shape identity became decodable shortly after stimuli appeared on screen, and extended to approximately 500 ms following their disappearance (Figure 3A, solid green bar indicates significant cluster). After the signal dropped to baseline levels, the presentation of the salient impulse revived decoding for a brief period of time (Figure 3B, solid green line indicates significant cluster). The averaged Mahalanobis distances showed a gradual reduction in pattern similarity for increasingly dissimilar shapes, suggesting a parametric relationship.

**Figure 3.**
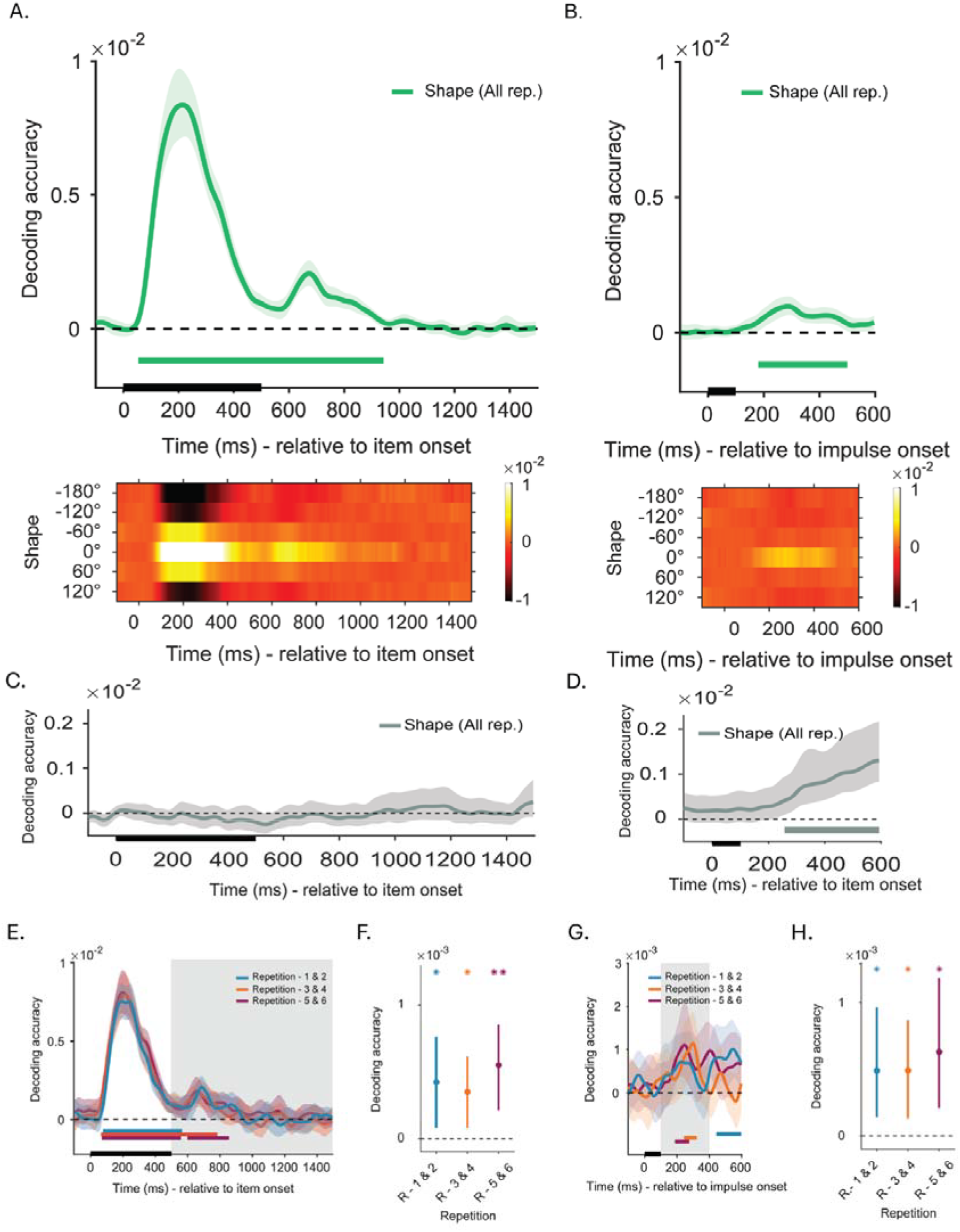
Decoding Results. **Note**: Time-course decoding of shape stimuli, relative the presentation of the memory item **(A)** and impulse signal **(B)** with respective similarity matrices presented at lower panels. The gradual change in distance as a function of angular difference between shapes suggest a parametric relationship between shape stimuli. (**C)** Time-course decoding of shape stimuli using gaze position recorded by eye tracker relative to memory item onset and **(D)** relative to impulse presentation. **(E)** Time-course decoding of shapes in Repetition 1&2 (blue), Repetition 3&4 (yellow) and Repetition 5&6 (red) bin relative to item onset and **(G)** impulse onset. **(F)** Mean decoding accuracy in each repetition condition within 500-1500 ms window relative to item onset and (**H)** 100-500 ms relative to impulse onset. **(A-B-C-D-E-G)** Solid lines represent mean decoding accuracy, the shadings mark 95% C.I., whereas solid bars at the bottom mark statistically significant time window. **(F&H)** Dots mark average decoding with error bars indicating 95 % C.I. Colored asterisks reflect average decoding above zero with statistical significance (* = *p* < 0.05; ** = *p* < 0.1).

Importantly, following concerns about the influence of eye movements on multivariate decoding of orientation gratings (Mostert et al., 2018) we also examined whether our stimulus space could be decoded from the eye position recorded by our eye tracker alone. We observed no significant clusters of decoding when shapes were encoded and retained (Figure 3C) providing little evidence that the peak EEG decoding observed during stimulus presentation and the subsequent 500 ms post stimulus offset was driven by ocular behavior. However, we did observe a significant cluster post-impulse as the response approached (Figure 3D) emerging after our impulse driven EEG decoding (approximately 80 ms difference in significant cluster onset times) and ramping up in anticipation of the memory probe presentation, which occurred 600 ms after impulse onset.

To assess the effect of repetition, we grouped each pair of subsequent repetition conditions (Repetition 1&2, Repetition 3&4 and Repetition 5&6, see methods), randomly sampled trials to ensure equal distribution of shapes in each condition and repeated decoding analyses. The trial-specific shape identity could be reliably traced between approximately 100 - 400 ms window following memory item offset (Figure 3E). Assessing differences between repetition conditions against zero yielded no statistically significant clusters. Average decoding within 500 – 1500 ms period (corresponding to CDA window) was also above zero for each repetition condition according to group permutation test (Figure 3F; Repetition 1&2, blue, t(29) = 2.43, *p* = 0.022; Repetition 3&4, orange, t(29) = 2.5, *p* = 0.018; Repetition 5&6, red, t(29) = 3.45, *p* = 0002). The repetition conditions did not differ from each other according to an RM-ANOVA with repetition as a main effect (*F*(2, 58)= 0.433, *p* = 0.65).

Next, the effect of repetition on trial specific shape identity decoding was assessed in post-impulse period. We observed statistically significant clusters in all repetition conditions (Figure 3G), and the temporal average within 100-500 ms period relative to impulse onset was above zero in all conditions (Figure 3H, Repetition 1&2,, t(29) = 2.7, *p* = 0.012; Repetition 3&4, t(29) = 2.85, *p* = 0.009; Repetition 5&6, t(29) = 3.3, *p* = 0.003). The repetition condition did not yield any differences in decoding accuracy when temporal averages were compared via an RM-ANOVA with repetition as a main effect (*F*(2, 58)= 0.29, *p* = 0.75).

## 3. Discussion

Extensive work in cognitive psychology has demonstrated the beneficial effect of repetition on memory (Ebbinghaus, 1913). Corresponding encephalographic work has shown that repetition leads to a reduction in the CDA, an index of memory load, and this reduction has been taken as evidence for a representational shift to LTM. However, the CDA is thought to represent an episodic marker or pointer towards the memorized information rather than representing the content itself. Here we more directly investigated the effects of repetition on mnemonic content by adopting a multivariate decoding approach in combination with a continuous shape space (Li et al., 2020). Importantly, we observed a clear dissociation between the CDA and content-specific decoding performance measures as a function of repetition. Consistent with earlier studies (Carlisle et al., 2011; Gunseli, Meeter, et al., 2014; Gunseli, Olivers, et al., 2014; Özdemir et al., 2024; Şentürk et al., 2024; Woodman et al., 2013), we observed a decrement in the CDA across repeated presentations of the memory item. In contrast, shape decoding showed no such decrement and remained constant across repetitions – if anything, numerically, shape decoding became more reliable for the later repetitions (though this was not born out in the statistics).

The successful decoding of shape from VWM adds another useful feature dimension to the visual properties that have been found to be decodable from the EEG (i.e. orientation and color; Hajonides et al., 2021; Kandemir, Wilhelm, et al., 2024; Wolff et al., 2015). Moreover, during the initial presentation and retention interval, we could not extract the shape information from the eye coordinates, making it unlikely that the activity pattern was an artifact derived from eye movements (Mostert et al., 2018). However, following impulse presentation and immediately before the test item appeared on the screen, we did observe a reliable ramp-up in ocular derived decoding of current shape memory item. These results seem likely to indicate learned preparatory ocular behavior meant to shift attention towards locations where targets and foils were known to differ the greatest, thereby aiding in the detection of foils. Overall, these results demonstrate reliable decoding of Li et al.’s (2020) continuous shape space (see also: Printzlau et al., 2026). However, there the decoding observed immediately before the probe stimuli were presented on the screen suggests some strategic ocular contribution, likely displacing the eye towards the spatial location known to contain the largest difference between individual foils and test stimuli learned through experience. As such, some caution is called for in future experiments utilizing a continuous shape space with multivariate decoding, as ocular contributions are likely in the period immediately before test stimulus presentation.

We observed stable decoding across repetitions during the same time window as the CDA, as well as for the re-emerging pattern of activity following the perturbation stimulus (“ping”). If we take decoding performance as a measure reflecting content-related neural representations, then the conclusion is that these representations did not weaken with repetition and hence do not provide evidence for a shift towards LTM. Moreover, the observed dissociation supports the idea that the CDA does not reflect content activity, but instead reflects pointer-like information providing spatial or object-based embedding of the memory (Balaban, 2026; Balaban & Luria, 2015, 2025; Luria & Vogel, 2011; Styrkowiec et al., 2026). Such pointer information may initially be helpful in providing access to the memory, but with repetition, as the memory content becomes familiar, it becomes less useful. It is also important to point out that the fact that shape decoding remained stable across repetitions does not mean that shape was not learned: observers clearly became better on the memory task with repetition. What it would mean is that VWM retained the content representations during learning, at least across the limited range of trials explored here.

The claim that content representations remain active in VWM across repetitions primarily rests on the decodable activity patterns during the same time window as the CDA. However, as can be clearly observed also in our data, such decodable activity typically declines with time into the delay interval, raising the possibility that the memory transitions to an activity-silent, connectivity-based storage format already within each trial. Replicating earlier findings (Duncan et al., 2023; Kandemir, Wolff, et al., 2024; Wolff et al., 2017), we found that such patterns can then be recovered by a perturbation stimulus, and we extend these findings by showing that also these revived activity patterns remain stable across repetition (or if anything improve in terms of statistical reliability). The question then becomes whether this activity-silent representational format still reflects working memory or reflects already the first stage or instance (Logan, 1988, 1990) of long-term memory formation. This question is not easy to answer, partially because the definitions of working memory and long-term memory are functional in nature and do not necessarily map one-to-one onto specific neurological processes. While some argue that working memory is, by its very nature, defined by continuous neural activity (Constantinidis et al., 2018; Foster et al., 2024; Fuster & Alexander, 1971; Zipser et al., 1993), others have argued that much of what has been attributed to active working memory may instead be accomplished by shifts in synaptic connectivity over short time periods (Barak & Tsodyks, 2014; Mongillo et al., 2008; Olivers et al., 2011; Polyn & Woodman, 2025; Stokes, 2015; Trübutschek et al., 2017), while again others proposed that short term memory tasks are accomplished via a combination of active working memory systems and silent long-term memory systems (Awh & Vogel, 2020; Foster et al., 2024; Hakim et al., 2021; Lundqvist et al., 2018). Several features of ping induced decoding observed in past studies suggest that it may indeed reflect the starts of long-term memory formation as episodic traces are being built up. For instance, ping induced decoding has previously been shown to contain previous trial information (Duncan et al., 2023; Zhang & Lewis-Peacock, 2024), tracks the order in which memoranda were encoded (Huang et al., 2021), is improved when the pings themselves match some aspects of the memory items (Karabay et al., 2025; Yang et al., 2025), generalizes to associated memoranda (Kandemir & Akyürek, 2023; Wolff, Kandemir, et al., 2020; but see van Bree et al., 2024), and that under certain conditions, allows decoding of temporarily accessory items within working memory tasks (Kandemir, Wilhelm, et al., 2024). Recent monkey physiology work has additionally shown that pings can decode both memorized stimuli, as well as passively viewed recently presented stimuli (Yiling et al., 2024). Each of these characteristics may to some degree be predicted by an episodic trace account of impulse evoked decoding where latent traces in the memory system passively build up and interact with one another (Beukers et al., 2021). In the current study we found that ping-induced decoding was relatively durable across repetitions, suggesting impulse decoding is mostly sensitive to the most recent memory trace with little sensitivity to trace accumulation.

One possible explanation for the familiar CDA decrement (that is consistent with the proposal that the CDA reflects object pointers) is that the memory changes across repetitions from location-specific formats to spatially generalized (Ester et al., 2009; Fukuda et al., 2016; Kandemir & Olivers, 2024, 2026) or potentially centralized formats (Chambers et al., 2013; Williams et al., 2008). With every repeated trial, the original location becomes less informative as a retrieval cue (since on preceding trials it may have been at the other location; but see Reinhart et al., 2014 for an example where CDA declines when locations are also repeated), with location information becoming less needed given that the content is being learned, and, ultimately, it is only the content that is relevant to the memory task. Thus, the decreasing CDA may not so much reflect offloading of the VWM to LTM, but changing the internal spatial mapping of the memory. To test for shifts in memory load, newly developed multivariate measures of working memory load may be more powerful as they do not require lateralized stimuli (Adam et al., 2020; Suplica et al., 2025; Thyer et al., 2022). Interestingly, this measure appears more sustained than the CDA during the retention interval that the CDA shows, suggesting that working memory load signals may persist even when the CDA disappears. Unfortunately, the current data falls short in this regard as we did not manipulate memory set size, and thus how this global measure of load changes with repetition remains a question for the future.

In sum, the exact mechanisms of learning and how they affect working memory representations remain elusive. What we have been able to demonstrate is a clear dissociation between CDA amplitude and stimulus-specific multivariate decoding as a function of repetition. This provides further evidence that the CDA does not reflect content activity per se, and that in this initial phase of learning what might primarily change is the specific contextual information (memory pointers) rather than memory content.

